# Photomap of the polythenic chromosomes of *Drosophila malerkotliana* and *in situ* mapping of the Hsp83 *locus*

**DOI:** 10.1101/2020.08.05.238600

**Authors:** Pierre Teodósio Félix, José Ferreira dos Santos

**Affiliations:** Laboratory of Population Genetics and Computational Evolutionary Biology - LaBECom, UNIVISA, Vitória de Santo Antão, Pernambuco, Brazil; Laboratório de Genética de Microrganismos Departamento de Genética, Universidade Federal de Pernambuco. Av. Morais Rego S/N. Cidade universitária, 50732-970, Recife (PE), Brasil

**Author notes:** Corresponding author/ **Contact**.

**Keywords:** *Drosophila malerkotliana*, Hsp83, Polythenic chromosomes, Photomap, *in situ* hybridization

## Abstract

A photomap of the polythenic chromosomes of *D. malerkotliana* was constructed to facilitate the identification of the chromosomal arms and their sections, in order to allow the identification of the breaking points of inversions and the location of the bands marked by *in situ* hybridization. The photomap included the six chromosome arms corresponding to pairs I (chromosome X), II and III, excluding pair IV and Y chromosome, because they were not visualized in the material examined, probably due to their small size. Through the *in situ* hybridization technique with the use of a biotined probe of a fragment of the *D. melanogaster* gene, the Hsp83 locus of *D. malerkotliana* was mapped. The probe hybridized with a frequency of 70% in section 98 of the IIIR chromosome. This is the first mapped gene in the species, and indicates that possibly the IIIR arm of *D. malerkotliana* corresponds to the IIIL arm of *D. melanogaster*, where the Hsp83 gene was located.

## Introdution

The mapping of genes by *in situ* hybridization is an excellent method to establish the homology of the polythenic chromosomes of species close to Drosophila, allowing to clarify their evolutionary relationships. The genes of thermal shock proteins (HSP), due to their high evolutionary conservation, are especially suitable for this approach, as they also allow comparison with other groups of more distant species. The Hsp70 gene of *Drosophila melanogaster* was used by BONORINO et al (1993) to map the corresponding locus in several species of the *willistoni* group, demonstrating hybridization in a single section of chromosome III of each species, and confirming the homology of this chromosome with the IIIR arm of D. melanogaster. The homology of the chromosomal arms between the two groups was extended by mapping the genes *Hsp83, Hsr, Hsp27* and *Ubi* (among others) of *D. melanogaster* in several species of the *willistoni* group, by RIEGER (1999). Similar studies had already been carried out for the obscura group (SEGARRA et al., 1999) and for the group full (RUIZ et al., 1997) of Drosophila species.

In this work we begin the study of the homology of the chromosomal arms of *D. malerkoltiana*, (subgroup *ananassae*) with *D. melanogaster*, (*melanogaster* subgroup) both of the *melanogaster* group, by mapping the Hsp83 gene. To facilitate the study it was nescessary to produce a photomap of the main chromosomal arms of *D. malerkotliana*, since the map drawn by JHA and RAHMAN (1973) and reproduced by SORSA (1988), proved to be difficult to compare with the chromosomes of the populations of the species collected in northeastern Brazil.

## MATERIAL AND METHODS

### Making the photomap of polythenic chromosomes

The slides of politenic chromosomes of *D. malerkotliana* of the population of Parque Dois Irmãos (Recife) were prepared by crushing the dissected salivary glands of zero-hour pre pupae, fixed with a solution of acetic acid, water and lactic acid, in a 3:2:1 ratio. The material was stained with aceto-lactic orcein (1g of orcein-MERCK, 45ml of acetic acid, 25ml of lactic acid 85%, and 30ml of distilled water), according to ASHBURNER (1967). The best slides were photographed under Leica Microscope in increase of 100 X 1.25 with Kodak TMAX Iso 100 film.

#### In situ hibridization

For *in situ* hybridization, the ENGELS et al. (1986) technique was used. After dissection of the larvae, the salivary glands were fixed in 45% acetic acid and crushed with cover slip in a lactic-acetic solution (lactic acid, water and acetic acid, 1:2:3). The slides were stored at 4°C for at least 18 hours. The slides were removed and the slides were submitted to several washes in 2X SSC and ethanol. The slides were air dried and stored at 4°C. After these steps, the best slides were selected and only those with intact chromosomes, spread properly, were stored for further hybridization.

For the marking and mapping of the Hsp83 locus in *D. malerkoltiana*, the clone λ6 of the Hsp83 gene of *D. melanogaster* (HOLMGREN et al., 1981), inserted in the plasmid pBR322, was used as probe. The plasmid was amplified by the transformation of the Xm1 strain of the bacterium *Escherichia coli*, cultivated in LB medium with ampicillin (100°g/ml) (SAMBROOK et al., 1989), and was extracted by non-phenolic method (PHILIPPSEN et al., 1991).

To prepare the probe, 1μg of the plasmid containing the gene fragment, marked with biotin by *nick-translation* with gibco’s Bionick kit, was used. The probe marking was tested in “*dot blot*” with revelation with Estreptavidin – Alkaline Phosphatase (SA-AP), Nitroblue tetrazolium chloride (NBT), 5-bromine-4-chloro-3-indolylphosphate p-Toluidine salt (BCIP). 500ng of probe per blade was used under high astringency conditions (temperature of 36° C and 50% of formamide) for 48 hours. The revelation was performed with SA-AP, NBT and BCIP, followed by the lacto-acetic orcein counter-dye at 0.2%. After drying the blades, the material was permanently assembled with Entellan.

## Results

The photomap of the polythenic chromosomes of *D. malerkotliana* was constructed with the six chromosome arms corresponding to pairs I (chromosome X), II and III, which are submetacentric (Figure 1). Pair IV and Y chromosome were not included because they were not visualized in the material examined, probably due to their small size. The photomap is essential to facilitate the identification of the chromosomal arms and their sections, in order to allow the definition of the breaking points of inversions and the location of the bands marked by *in situ* hybridization. Whenever possible, the sections of the photomap were marked according to the polythenic map drawn by JHA and RAHMAN (1973) for the species, reproduced by SORSA (1988). Additionally, the largest sections were divided into subsections.

**Figure 1.**
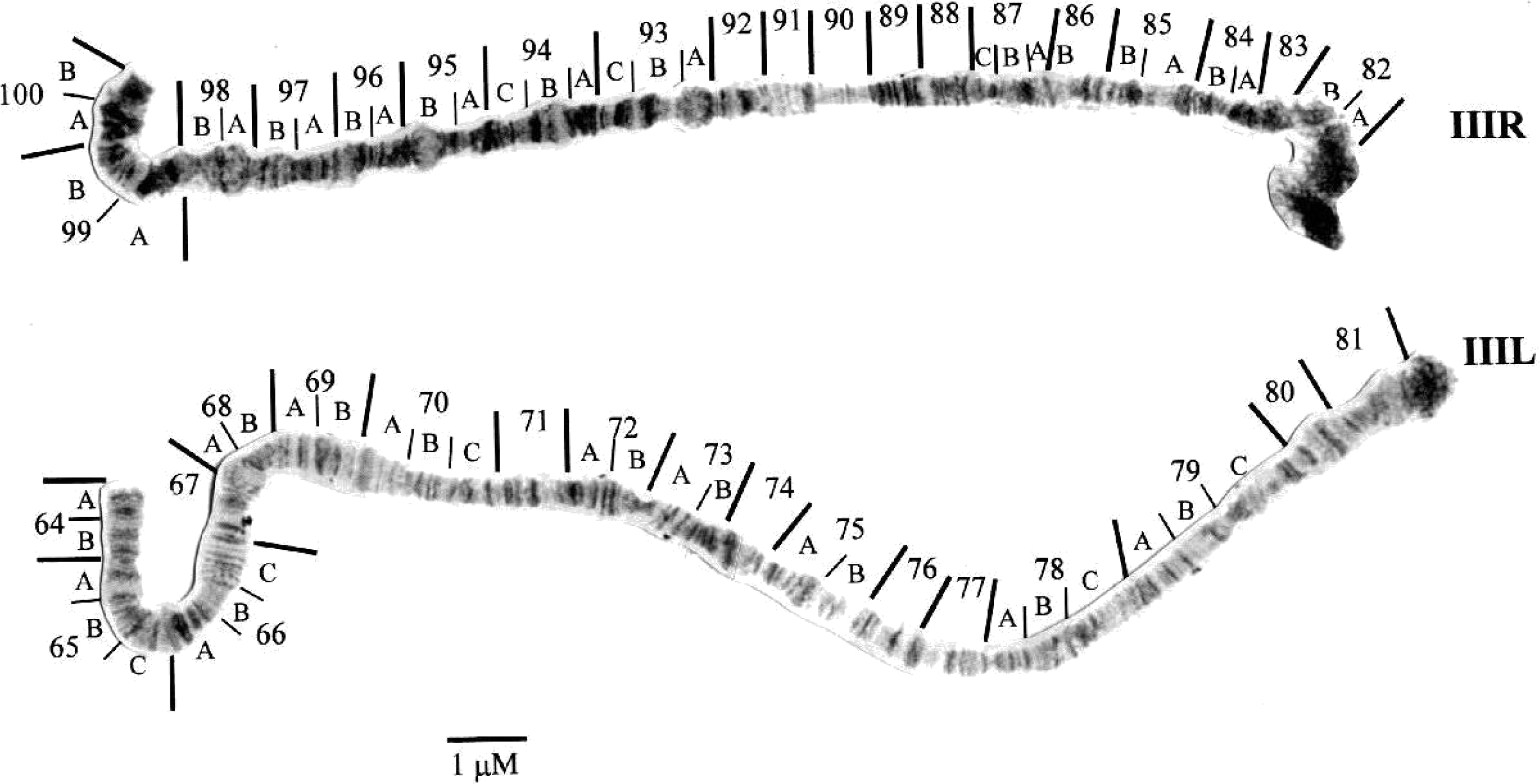
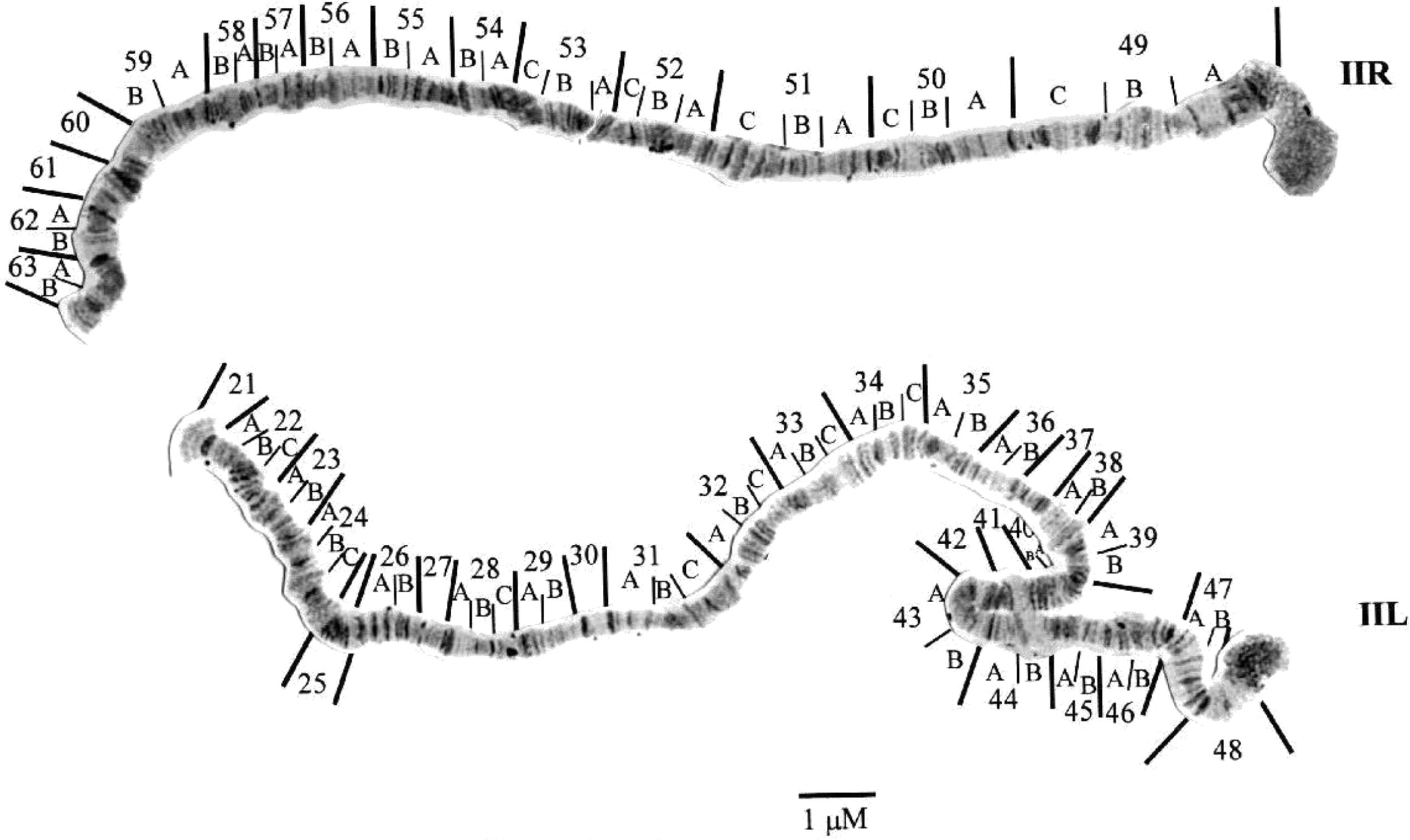
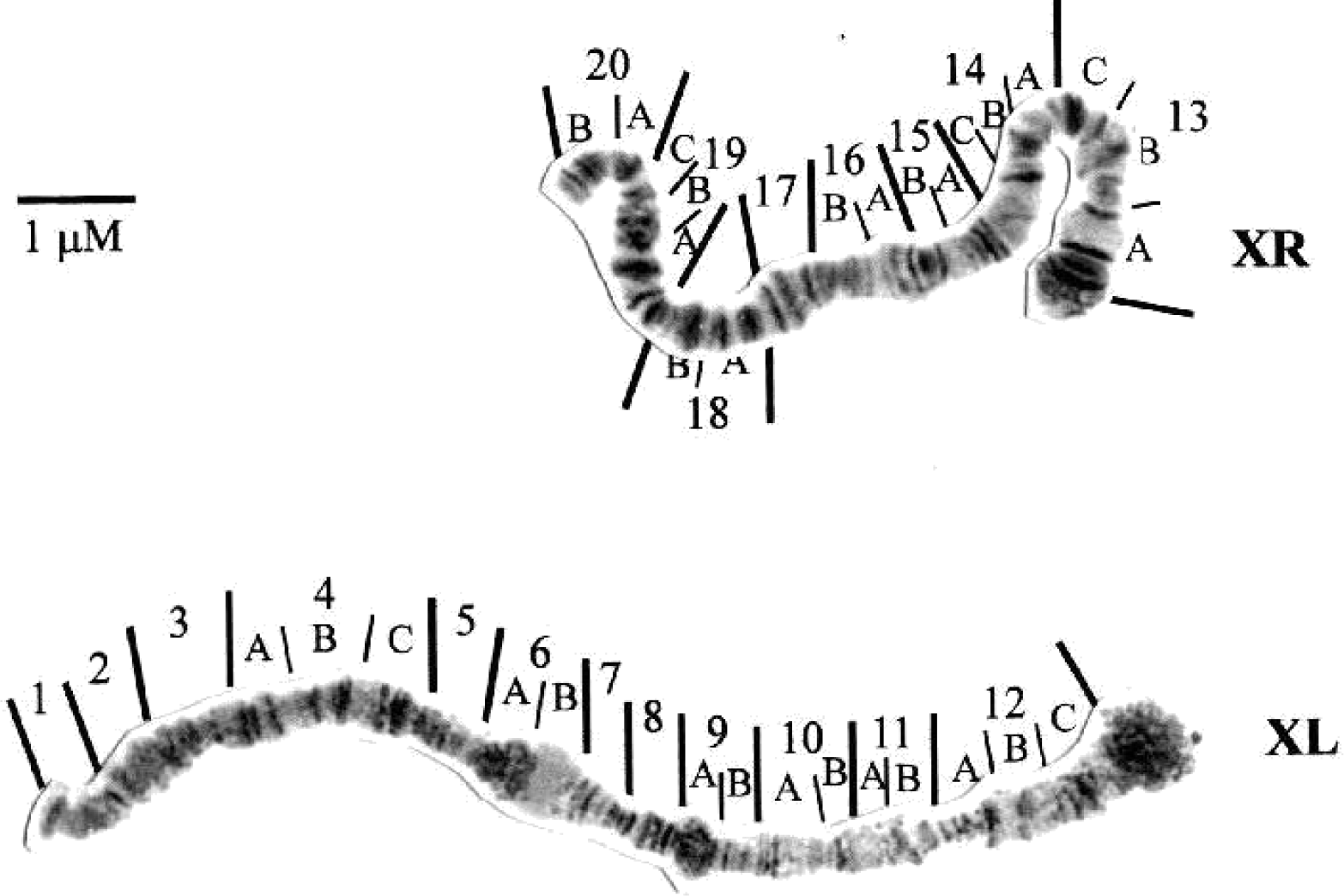
Photomap of the polythenic chromosomes of *D. malerkotliana*. The tip (telomere) of each chromosomal arm is positioned on the left and the base (centromere) on the right. The division of sections followed the map drawn by JHA and RAHMAN (1973) and reproduced by SORSA (1988) for this specie. In this photomap, each section was subdivided into subsections for easy use. Range: Bar =12.6mm = 1μm

*In situ* hybridization was performed on polythenic chromosome slides of 3 individuals. Fourteen marks were observed in section 98 of the chromosomal arm IIIR (Figure 2), representing 70% of the markings. Other marks were found in sections 91 (2 marks, 10%), 87 (2 marks, 10%) and 85 (1 mark, 5%) of the same arm and in section 25 (1 mark, 5%) of the IIL chromosomal arm.

**Figure 2.**
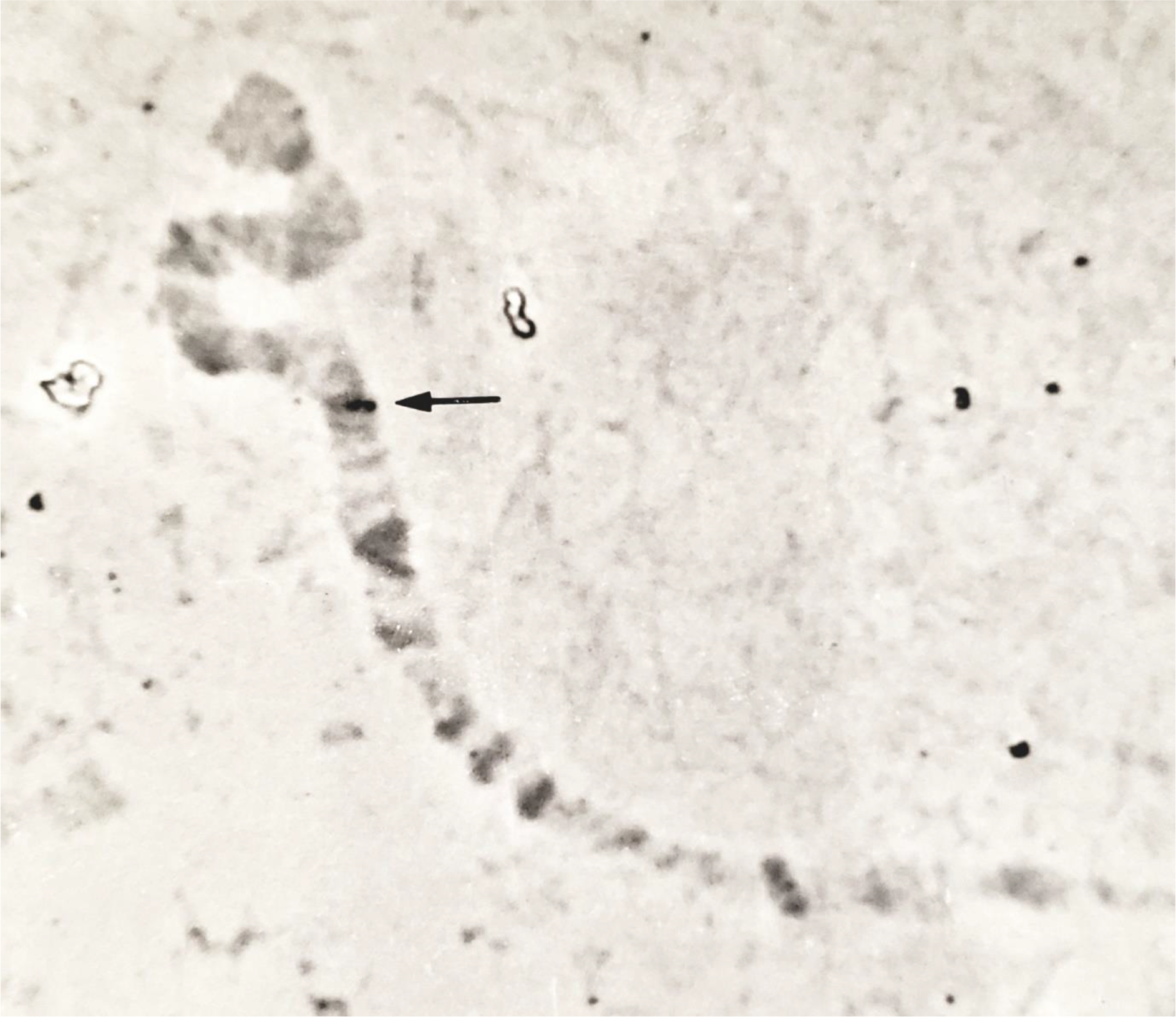
Tip of the chromosomal arm IIIR of *D. malerkoltiana*, showing the hybridization of the Hsp83 gene probe in section 98. This marking was found in 14 nuclei, representing 70% of the total markings observed.

## Discussion

The nuclei of *D. malerkotliana* presented six chromosomal arms, similar to the other species of the *ananassae* subgroup and different from *D. melanogaster*, from the melanogaster group, which has five chromosomal arms (LEMEUNIER et al., 1986). The six arms correspond to pairs I (chromosome X), II and III. The PAIR IV and the Y chromosome, although included in the map drawn by JHA and RAHMAN (1973), were not visualized in the analyzed material, possibly due to its small size and its non-polythenization, which is common in several species of *Drosophila* (SORSA, 1988). The map of JHA and RAHMAN (1973) presents several divergences of bands, inter-bands and “puffs” in relation to the polythenic chromosomes of the populations of the Northeast. This difference is probably related to changes in the chromosomal constitution, resulting from the accumulation of variations over the colonization time, possibly introduced by accumulated inversions and regulatory divergences, due to the large geographical distance from the distribution center of the species.

The Hsp83 gene was mapped in *D. malerkoltiana* in the section 98 of the chromosomal arm IIIR, with frequency of 70%, being the most consistent mark observed. Whereas in D. melanogaster the Hsp83 gene hybridizes in section 63BD of arm IIIL (ZHIMULEV et al., 1974), this result indicates probable homology of the chromosomal elements IIIR and IIIL between the two species. Additional studies including hybridization of other genes may confirm this homology.

The other hybridization marks, with less consistent results, probably represent other genes of the *Hsps* family, which present evolutionarily conserved regions (CRAIG et al., 1993). The hypotheses of the presence of pseudogenes or duplication are less likely, as occurs for the *Hsp70* locus in *D. melanogaster*, occupying sections 87A and 87C of the IIIR arm (LIVAK et al., 1978; VAZQUEZ et al., 1993) and for the *Hsp83* locus in *D. insularis*, which has two equally marked regions, one on the XR chromosome (as in any *willistoni* group) and the other in the IIR that is exclusive to it (RIEGER, 1999).

In this work, a photomap was made and mapped by *in situ* hybridization a physiologically important gene, the first being mapped in *D. malerkotliana*, representing the beginning of a complete study of the chromosomal evolution of the species, which will also include the induction of poufs by stress or hormones such as ecdysone.

## Attachments

**Figure 1.**
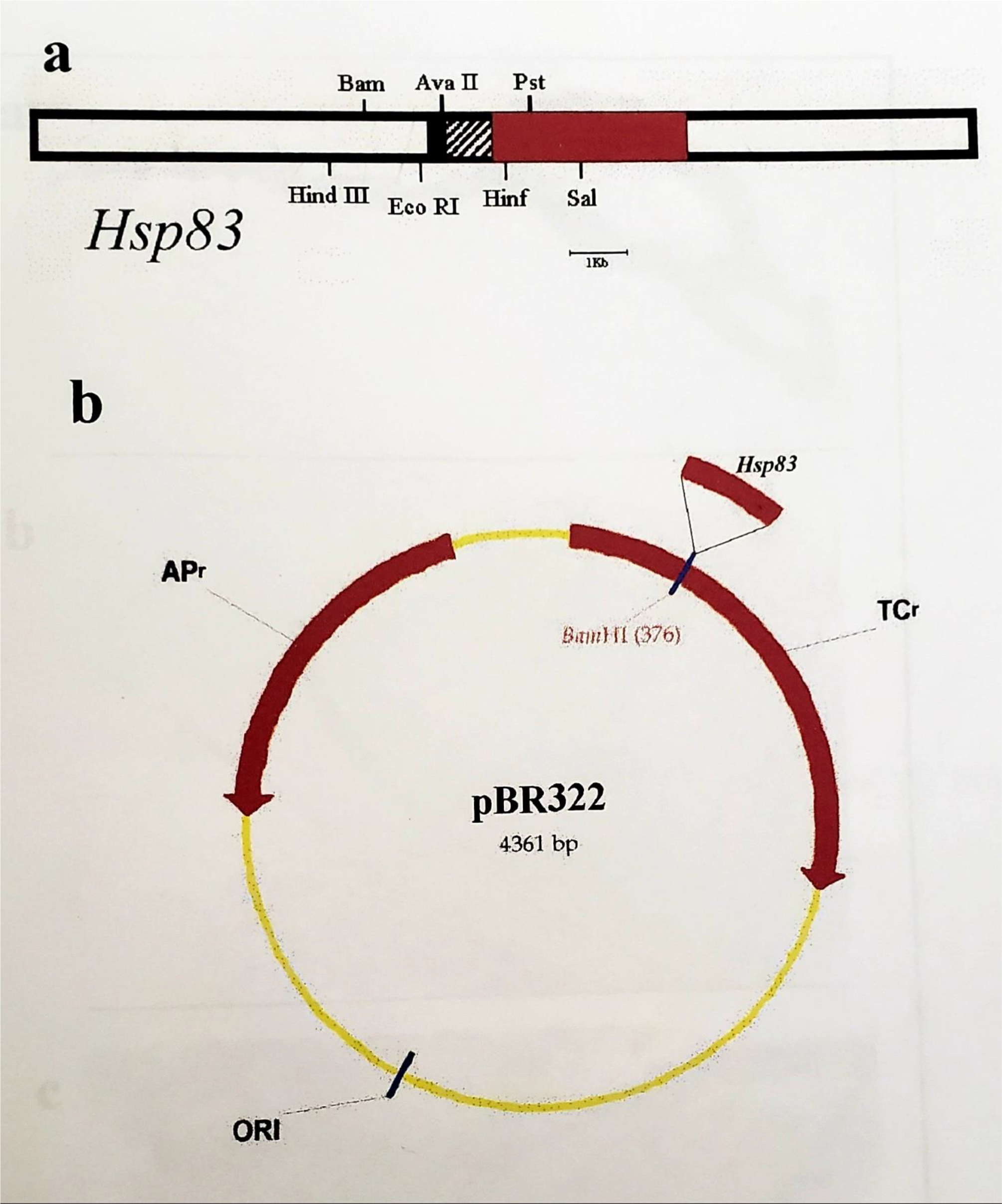
Restriction map of clone λ 6 with 3.4kb (a), containing the fragment of the Hsp83 gene of *Drosophila melanogaster* inserted in plasmid pBR322 (b) sequence homologies in the 5’ regions of four *Drosophila* heat shock genes (HOLMGREN and Cols., 1981).

